# ProteinGCN: Protein model quality assessment using Graph Convolutional Networks

**DOI:** 10.1101/2020.04.06.028266

**Authors:** Soumya Sanyal, Ivan Anishchenko, Anirudh Dagar, David Baker, Partha Talukdar

## Abstract

Blind estimation of local (per-residue) and global (for the whole structure) accuracies in protein structure models is an essential step in many protein modeling applications. With the recent developments in deep-learning, single-model quality assessment methods have been also advanced, primarily through the use of 2D and 3D convolutional deep neural networks. Here we explore an alternative approach and train a graph convolutional network with nodes representing protein atoms and edges connecting spatially adjacent atom pairs on the dataset Rosetta-300k which contains a set of 300k conformations from 2,897 proteins. We show that our proposed architecture, ProteinGCN, is capable of predicting both local and global accuracies in protein models at state-of-the-art levels. Further, the number of free parameters in ProteinGCN is almost 1-2 orders of magnitude smaller compared to the 3D convolutional networks proposed earlier. We provide the source code of our work to encourage reproducible research.^1^

## 1 Introduction

Despite progress of deep learning-based methods as recently demonstrated in the CASP13 experiment^2^, protein structure prediction remains a challenging problem. This is especially true if one needs models of atomic-level accuracy - the resolution required for biologically relevant applications such as, molecular replacement, small-molecule binding or protein-protein interaction studies. Along with the conformational space sampling, the scoring function is the key component of modeling, which allows for proper ranking of the putative models and selection of the ones closest to the native structure. Estimating both global and local per-residue scores are important. The former provides an overall perspective on model’s quality. The latter differentiates between regions within a model with respect to the degree of their structural similarity to corresponding local regions in the native structure. This is especially useful for subsequent protein structure refinement.

Various methods have been developed to tackle the scoring problem, ranging from energy functions derived from either general physical principles (like a number of popular molecular mechanics force fields CHARMM [18], AMBER [6], OPLS [13], GROMOS [30]), or deduced from diverse sets of known protein structures (various knowledge-based or statistical potentials GOAP [38], RW [36], DFIRE [39]), or both (Rosetta [17]), to the ones trained to estimate particular similarity scores (e.g., lDDT [20], CAD [22], GDT [35], TMscore [37], etc.) between a computational model and the experimentally determined reference structure directly from atomic coordinates of the former. Meta-methods combining one or more of the above scores with additional biological data (e.g., multiple sequence alignments, templates, etc.) also exist; some of them use neural networks to learn the mapping of the above hand-crafted features to the target similarity score (e.g., ProQ3D [32]).

The unifying idea among most scoring methods is that only spatially adjacent atoms or residues contribute to the quality score - this is due to the general principle of locality of inter-atomic interactions. To tackle the problem of model quality estimation from deep learning perspective, one approach is to project the structure onto a 3D grid and use 3D convolutions (convolutions are local by definition) to convert this voxelized representation in the quality score (3DCNN [5]). Lack of rotational invariance of this representation can be partly mitigated by data augmentation (use multiple different orientations for every model) during training. More recently, in the Ornate method [24] every residue is placed into its own local reference frame and convolutions are performed independently over a fixed-size box around each residue.

Another arguably more natural way of representing a protein molecule is by a graph with nodes representing atoms, and edges joining atom pairs which are closer in the 3D model than some predetermined *D_max_*; this representation is rotationally invariant by construction. Recently, there has been much progress in the field of *Geometric deep learning* [2] which deals with developing graph based deep neural networks for graphical structures [10, 29, 16, 3]. Such methods have the ability to automatically learn the best representation (embedding) from raw data of atoms or bonds features for different node-level and graph-level attribute predictions. These approaches have been successfully applied to molecules for performing various tasks such as molecular feature extraction [7, 14, 9], drug discovery [1], protein structure and crystal property prediction [34, 28]. In most molecular GCNs, edges encode information on spatial proximity of atoms, but not on their mutual orientations. In this work, we explicitly take into account inter-atomic orientations, and extend the application of GCN to the protein model quality assessment problem. We show that, with 20-fold less learnable parameters than in 3D CNNs, the new method called ProteinGCN shows state-of-the-art performance on diverse benchmarks derived from CASP13 server submissions, as well as on extensive Rosetta decoys.

## 2 Background: Graph Convolutional Networks

In this section, we give a brief overview of Graph Convolutional Networks (GCNs) [16] for undirected graphs. Let 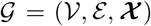 be a graph, where 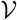 denotes the set of vertices, 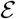 represents the set of edges, and 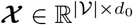 represents *d*_0_-dimensional input features of each node. A single GCN layer update equation to obtain the node representations is defined as:

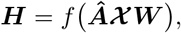

where, 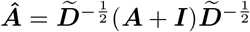 is the normalized adjacency matrix for a given adjacency matrix ***A*** after adding self-loops, and 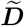 is defined as 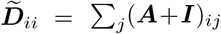. 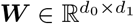 denotes the model parameters and *f* is some activation function. The GCN representation 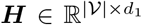 encodes the immediate neighborhood of each node in the graph. To capture multi-hop dependencies in the graph, multiple GCN layers can be stacked as follows:

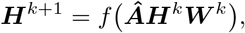

where *k* denotes the number of layers, 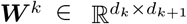 is layer-specific parameter and 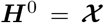. For a single vertex with representation ***υ_i_* ∈ *H***, the same equation can be written as follows:

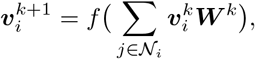

where 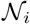 denotes the neighborhood of the *i^th^* node. We will use this vertex based update equation to describe our GCN formulations.

## 3 ProteinGCN

In this section, we first describe the problem statement in detail (Section 3.1). Next, we discuss the various steps involved in training ProteinGCN. In that, we first explain the process of generating input protein graph required for training ProteinGCN (Section 3.2). Then, we formally define the model architecture (Section 3.3). Finally, we specify the loss criteria used to train the ProteinGCN model (Section 3.4).

### 3.1 Problem Formulation

Protein model quality assessment (QA) is broadly defined as the task to score protein conformations such that the ones closest to the native structure can be identified using the predicted scores. It has two related sub tasks - predicting a global score for the protein conformation and predicting local scores for each amino acid residue in the protein.

### 3.2 Protein Graph

A natural way to represent any given protein structure is by modeling it as a graph. Such a construct has been shown to be effective in learning a representation of a local neighborhood around each amino acid residue [8]. In this work, we create a *protein graph* with nodes representing the various constituent non-hydrogen atoms in the protein. To connect the nodes in the protein graph, we consider *K* nearest neighbors for each node atom and connect the atom to each of the *K* neighbors. For computing the distance between atoms, we use the given 3D protein structure. Note that, with this formulation for graph connectivity, we can capture the 3D interactions in a better way since the connections are not restricted to only the chemical bonds present in the protein structure. Now, given this protein graph, we use one-hot vector encoding as features to represent the atoms as nodes. We treat each heavy atom in the 20 standard residue types separately, which yields 167 atom types and hence the dimension of the node feature vector. For edges, we explore three types of features as described below:

1. **Edge distance [ED]:** Every edge is assigned a distance measuring the proximity of the two atoms in the protein structure model. We use gaussian basis expansion to generate a vector representation for the distance. In this work, we use gaussian basis with means varying uniformly from [0, 15] with a step size of 0.4.
2. **Edge coordinates [EC]:** To account for apparent anisotropy in relative atomic placements inside a protein, we complement the above distances with a set of directional features. To do this, a local reference frame 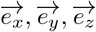 is placed at every atom, which is calculated from 3D coordinates of the two adjacent bonded atoms A,C and the atom in question B (see supplementary Table S1 for details):

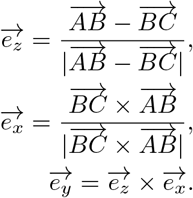 Every edge (*i, j*) then gets 2 × 3 additional features representing projections of atomic coordinates of atom *j*(*i*) onto the local frame of atom *i*(*j*).
3. **Bond type [BT]:** It is a binary feature which is 1 if the edge connection is an actual chemical bond in the protein structure and 0 otherwise.

### 3.3 ProteinGCN formulation

ProteinGCN utilizes graph convolutional networks (GCNs) to model the protein graph described in Sec. 3.2. Formally, let 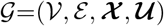 denote a protein graph. Here 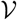 represents the set of atoms constituting the amino acid residues in the protein structure, 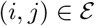 denotes an edge between atoms *i* and *j*, and 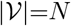 is the number of atoms in the protein graph. 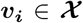 contains the features of the *i^th^* atom which can encode various properties of the atom. In this work we utilize one-hot vector for node representations. 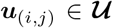 is the feature vector for the bond between atoms *i* and *j* that captures properties of the edges. To capture neighborhood influence using ProteinGCN, we utilize the graph convolution formulation as proposed by [34] for crystal graphs. It is defined as follows:

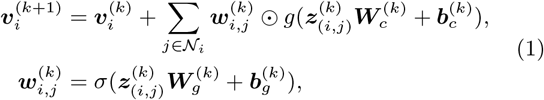

where 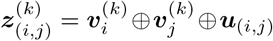 denotes the concatenation of atom and bond feature vectors of the neighbors of *i^th^* atom, ☉ denotes element-wise multiplication. *σ* denotes a sigmoid function and *g*(·) denotes any non-linear activation function (like ReLU). 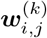 is an edge-gating mechanism [19] to incorporate different interaction strengths among neighbors. 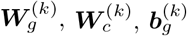 and 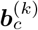 are the gate weight matrix, convolution weight matrix, gate bias, and convolution bias of the *k*-th layer of GCN respectively.

As discussed in Sec. 3.1, the task of protein QA involves predicting scores at the global protein level as well as at the local amino acid residue level. In other words, it can be seen as a regression problem at graph (protein graph) and subgraph (amino acid residue) level respectively. Graph pooling [11, 27] is a technique to generate a graph representation using the learnt node embeddings. A simple example of pooling is the average operator which takes the mean of the representations of nodes in the graph. Similar ideas can be extended to generate a subgraph embedding using its constituent nodes. Hence, in ProteinGCN, the learnt atom features are then averaged to get a vector representation of the protein graph. In a similar manner, representations for each amino acid residue are calculated by averaging the embeddings of the constituent nodes. More concretely,

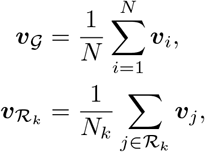

where 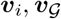, and 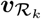 are the atom, graph and *k^th^* amino acid residue 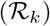 representations respectively. *N_k_* denotes the count of atoms that consitute the *k^th^* amino acid residue 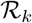.

Now, the global protein graph embedding 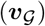 and local amino embeddings 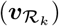 are fed to two separate fully-connected layers with non-linearities that learn to predict the global and local QA score respectively as defined below:

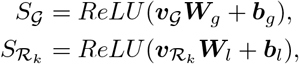

where 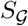 and 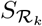 are the global and *k^th^* local scores respectively. ***W**_g_, **W**_l_, **b**_g_*, and ***b**_l_* are weights and biases for global and local fully-connected layer respectively.

### 3.4 Loss Criterion

We utilize the Mean Squared Error (MSE) as the objective function which measures the average squared difference between the estimated values and the actual target value for penalizing ProteinGCN while training the model.

Loss defined for the residue (local) score predictions is as follows:

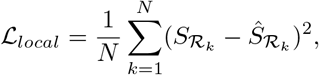

where 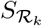 and 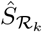 represent the *k^th^* residue (local) true score and the predicted score respectively. Similarly, the loss defined for global score predictions is as follows:

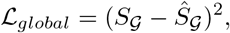

where 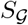 and 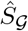 represent the global true score and the predicted score respectively for each protein. The combined loss 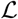 equally weights both local 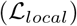 and global 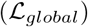 losses as follows:

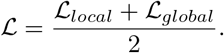

## 4 Experimental Setup

In this section, we provide details of the datasets and baselines used for the experiments, the evaluation strategy, and ProteinGCN hyperparameters.

### 4.1 Protein datasets

We use two protein datasets for all our experiments. Details about the dataset are provided below:

- **Rosetta-300k**: The primary set used for traing is composed of 2,897 protein chains ranging from 50 to 300 residues in length and having resolution not worth than 2.5A. For every protein chain, we generated 100 homology models of various accuracy using RosettaCM protocol [31] followed by dual-space relaxation [4] (more details on this set are given in Hiranuma et al [under submission]).
- **CASP13**: This set includes models (150 per target) for 80 target proteins from CASP13 experiment submitted by participating servers and selected by the organizers for the second stage of the EMA (Evaluation of Model Accuracy) experiment [33]. Similar to the Rosetta-300k dataset, all models were subject to dual-space relaxation in Rosetta to mitigate the possible diference in modeling procedures between different servers and to consolidate the mmodels with the ones from the training set.

### 4.2 Baselines

To compare the performance of ProteinGCN, we use the following baselines:

- **VoroMQA** [23]: It estimates protein quality by contsructing a Voronoi tesselation for the set of atoms in the protein model and then uses the derived inter-atomic contact areas to produce scores at atomic, residue and global levels.
- **Ornate** [24]: In Ornate, residue-wise lDDT scores are predicted from local 3D density maps by a deep 3D convolutional neural network. We have retrained the original network using the Rosetta-300k dataset to facilitate comparison with ProteinGCN.
- **ProteinGCN-Base**: This is a variant of ProteinGCN where we use only the edge coordinates [EC] as edge features. Further, we also restrict to using only the residue-level loss 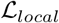. With these restrictions, ProteinGCN-Base is directly comparable to Ornate.

### 4.3 Evaluation

The scores during training are evaluated using Mean Absolute Error (MAE) accuracy metric. Finally, the Pearson correlation coefficient i.e., Pearson’s r is used to measure the linear relationship between the reference lDDT and the predicted lDDT scores for the Protein QA on the test set. This ultimately helps in understanding how close a model is to the native structure based on the predictions and the ground truth values. The Pearson’s r calculated for local residue predictions first involves calculating Pearson’s r for each local residue as shown below:

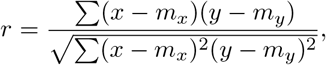

where *x* and *y* are the local residue prediction and target vectors respectively. *m_x_* is the mean of prediction vector *x* and *m_y_* is the mean of target vector *y*. We now average these local residue Pearson’s r coefficients over the total residues in a protein to get the local score as follows:

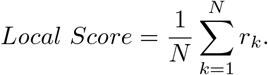

Similarly, to calculate Pearson’s r for the global predictions, we calculate Pearson’s correlation coefficient for each global value and finally take an average over all the coefficients.

## 5 Results

In this section, we evaluate the performance of ProteinGCN on multiple datasets and compare them with existing baselines. Next, we perform an ablation study to see the effect of different edge features on model’s performance.

### 5.1 Performance Comparisons

In order to evaluate the effectiveness of ProteinGCN, we compare it against the existing protein quality assessment baseline models listed in Section 4.2. The results are summarized in Table 1. We observe that ProteinGCN considerably outperforms all the baselines on both the datasets. Further, we find that ProteinGCN-Base shows consistent improvement over Ornate, even though both use the same set of features. The main difference between the two models is the use of GCN in ProteinGCN-Base compared to 3D-CNN in Ornate. This clearly demonstrates that GCN is better suited to model protein structures than the 3D-CNNs.

**Table 1:** Comparisons of ProteinGCN with baselines on two protein datasets. The pearson correlation is reported for all models. Please refer to Section 4.3 for evaluation strategy. Global correlation is missing for ProteinGCN-Base as it doesn’t have the global loss objective. We observe that ProteinGCN consistently improves upon previous baselines. For more details, please refer to Section 5.1.

| Method | Rosetta-300k |  | CASP13 |  |
| --- | --- | --- | --- | --- |
|  | Global | Local | Global | Local |
| VoroMQA | 0.738 | 0.478 | 0.785 | 0.387 |
| Ornate | 0.820 | 0.622 | 0.779 | 0.489 |
| PROTEINGCN-BASE | - | 0.674 | - | 0.548 |
| PROTEINGCN | **0.837** | **0.704** | **0.809** | **0.582** |

### 5.2 Ablation Study

To further evaluate the effect of various edge features and loss terms in the ProteinGCN model, we perform an ablation study over the two datasets described in Section 4.1. We sequentially remove some features from ProteinGCN model and evaluate its performance on the two datasets. The results are shown in Table 2. We observe that removing the global loss leads to a significant decrease in performance on both the datasets. Also, capturing the coordinate information leads to some gains in local prediction indicating the usefulness of capturing the edge orientations.

**Table 2:** Ablation of various components of ProteinGCN. The pearson correlation is reported for all models. 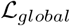 refers to global MSE loss used to train the model (Section 3.4). ED, EC, and BT are the edge distance, edge coordinate, and bond type respectively. Global correlation is marked with ‘-’ for models which do not have the global loss component. Please refer to Section 5.2 for more details.

| Ablation | Rosetta-300k |  | CASP13 |  |
| --- | --- | --- | --- | --- |
|  | Global | Local | Global | Local |
| **ProteinGCN** | 0.85 | 0.704 | 0.809 | 0.582 |
| **ProteinGCN** - EC | 0.85 | 0.7 | 0.787 | 0.571 |
| **ProteinGCN** - EC - BT | 0.851 | 0.701 | 0.787 | 0.575 |
| **ProteinGCN** - ED - BT - $\mathcal{L}_{global}$ | - | 0.674 | - | 0.548 |
| **ProteinGCN** - EC - BT - $\mathcal{L}_{global}$ | - | 0.641 | - | 0.525 |

### 5.3 Qualitative Analysis

To better understand the performance of ProteinGCN, we perform a quantitative analysis of the model’s prediction for a sample protein target (T1008). This is depicted in Figure 3. As shown in the Figure 2B, there is a strong correlation between the true and predicted global scores, which is desirable in real-case modeling scenarios when picking the best model from a pool of decoys is often a challenge. We also check how well local scores are recapitalated by picking three models at three distinct global accuracy levels and comparing predicted perresidue lDDT scores (shown in color) with reference ones (shown in grey, Figure 2C). Despite the differences in the global scores, ProteinGCN correctly captures trends in the local scores; this allows for selecting most inaccurate regions in the protein model which need further refinement [25].

**Figure 1:**
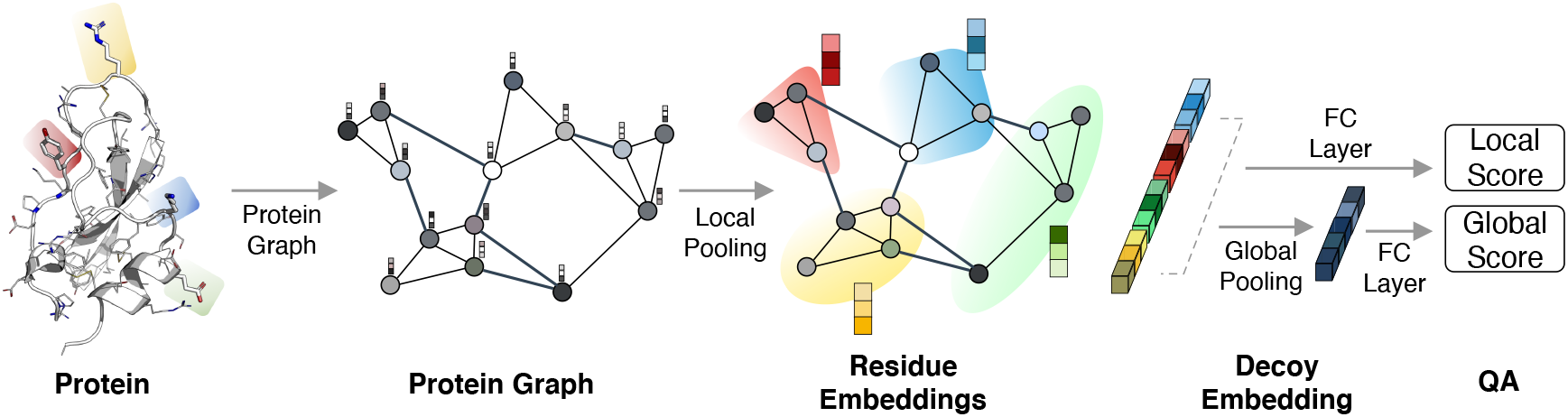
Overview of ProteinGCN: Given a protein structure, it first generates a protein graph (Section 3.2) and uses GCN to learn the atom embeddings. Then, it pools the atom embeddings to generate residue-level embeddings. The residue embeddings are passed through a non-linear fully connected layer to predict the local scores. Further, the residue embeddings are pooled to generate a global protein embedding. Similar to residue embedding, this is used to predict the global score. Please refer to Section 3.3 for more details.

**Figure 2:**
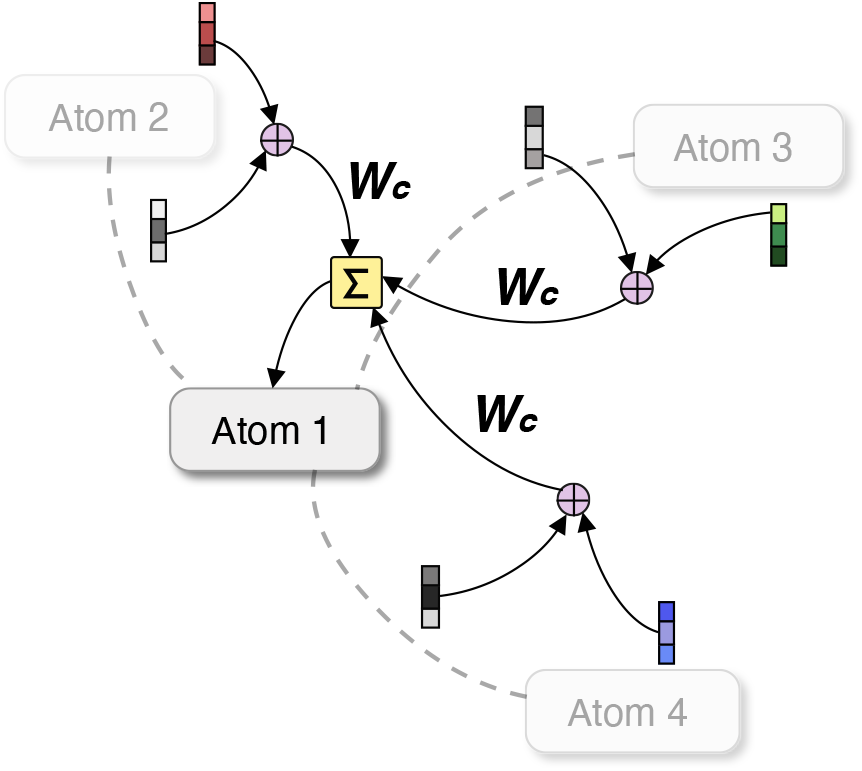
Overview of ProteinGCN update for a single node: For a given central protein atom node (e.g., *Atom 1* above) in the protein graph, ProteinGCN concatenates the neighbor embedding ***υ**_j_*, edge embedding ***u***_(*i, j*)_, and self embedding ***υ**_i_* using the concatenation operator ⨁. We omit concatenation of the self embedding in the diagram for clarity. The concatenated embeddings are then convolved with convolutional filter ***W**_c_*. The message from all the neighbors are then aggregated to get an updated embedding of the central node. The edgegating mechanism is also omitted in the figure for clarity (i.e., assume ***w**_ij_* = 1). Please refer to Equation 1 for more details.

**Figure 3:**
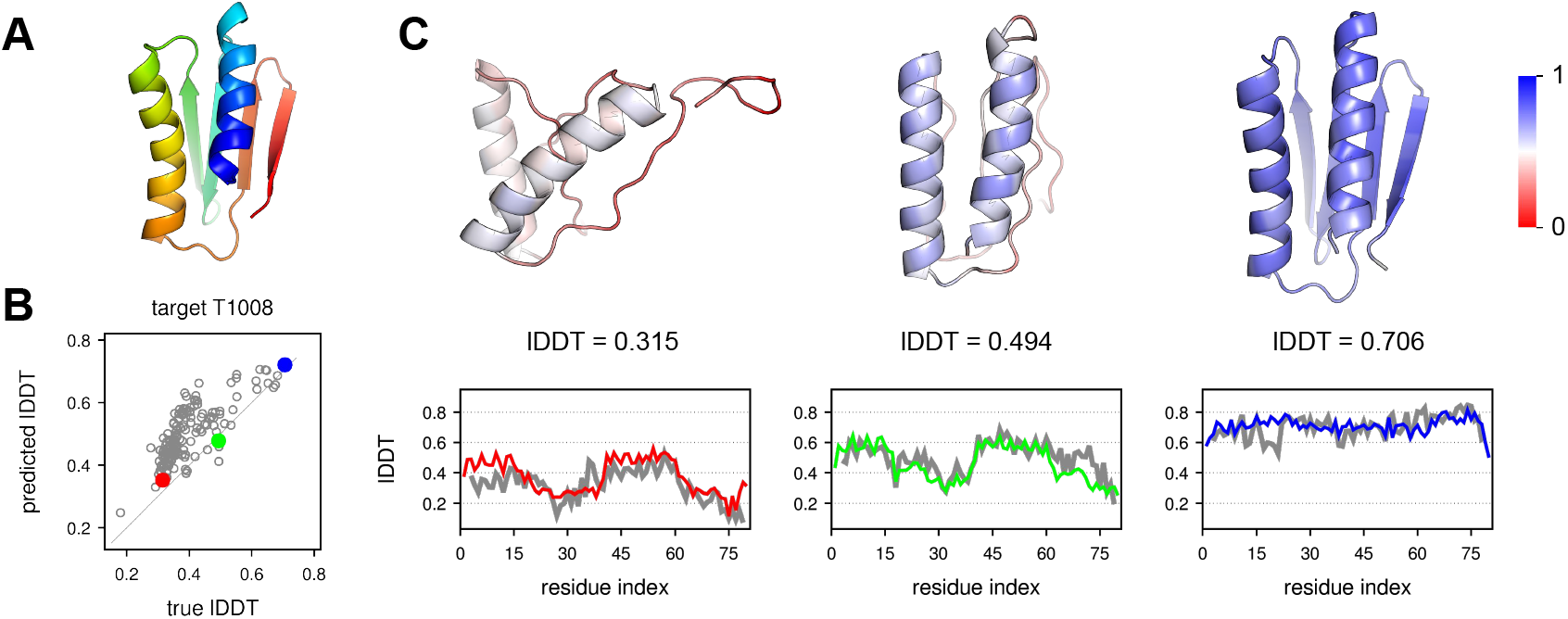
Qualitative analysis of the performance of ProteinGCN on CASP13 target T1008. (A) Experimental structure is shown in rainbow. (B) Predicted global lDDT scores are well-correlated with the reference scores with Pearson’s r = 0.778. (C) Three models of various accuracy levels (global lDDT scores are given on the Figure) are color-coded according to local lDDT scores predicted by ProteinGCN. Both reference (grey) and predicted (red, green, and blue) local scores are shown on the plots below the structures. We observe that ProteinGCN is able to correctly capture trends in local scores for different models of the target protein T1008. Please refer to Section 5.3 for more details.

## 6 Conclusion

In this work, we proposed ProteinGCN, the first graph neural network framework for the task of protein model quality assessment. Along with capturing the local structural information through the graph convolution formulation, ProteinGCN is also able to utilize both inter-atomic orientations and distances effectively. Along with that, ProteinGCN also utilizes 20x lesser learnable network parameters compared to Ornate, the state-of-the-art baseline. Through extensive experiments on two datasets, we establish the superiority of our proposed method over previous baselines.

## Acknowledgements

We would like to thank Hahnbeom Park and Nao Hiranuma for their constructive comments. This work is supported in part by the Ministry of Human Resource Development (Government of India).

## A Appendix

## A.1 Training Strategy

We first randomly split our dataset into 60/20/20 ratio of train, validation and test sets, unless specified otherwise. We then use one hot encoding to initialize the embeddings for all the 167 different protein atoms. The number of neighbours for each atom is set to 50, a maximum limiting value decided based on the closeness calculated using euclidean distance. Model weights are learnt and it converges after training ProteinGCN over 50 epochs using Stochastic Gradient Descent optimizer with the learning rate, lr=0.001 and the momentum parameter, m=0.9. To prevent overfitting and to regularize the model, we use Batch Normalization[12] within the convolution layer. Also, we use ReLU[21] as the activation function.

The model learns using the training set and then we check the test error on the validation set. We perform hyperparameter tuning over the hyperparameter space mentioned in the Table 3.

**Table 3:** List of hyperparameters and the ranges used for fine-tuning ProteinGCN model. Refer to Section A.1 for more details on training.

| Hyperparameter | Values |
| --- | --- |
| Number of convolutional layers, $n_{conv}$ | 1, 2, 3, 4, 5 |
| Hidden atom embedding dimension, $h_a$ | 64, 128, 512 |
| Hidden graph embedding dimension, $h_g$ | 32, 64, 128 |
| Number of fully connected layers, $n_{fc}$ | 2, 4 |
| Step size of the Adam optimizer, $lr$ | $10^{-4}$, $10^{-3}$, $10^{-2}$, $10^{-1}$ |

## A.2 Model Parallelization

ProteinGCN typically has around 100k trainable parameters (can change depending on the setting). The size of the model creates the process of batching a challenge and doesn’t allow larger mini-batches to fit in a single device for model training. Thus, we incorporated PyTorch [26] Data Parallel^3^ which splits and distributes the mini-batches across multiple GPUs evenly for the devices specified. This not only solved the problem of small MiniBatch sizes but helped us achieve performance gains in terms of 30% speedup while training on a 56 core machine configured with 6 NVIDIA GTX 1080Ti GPUs.

## Footnotes

1 https://github.com/malllabiisc/ProteinGCN

2 http://predictioncenter.org/casp13/

3 https://pytorch.org/docs/stable/_modules/torch/nn/parallel/data_parallel.html

