## Supplementary Table S1 for "ProteinGCN: Protein model quality assessment using Graph Convolutional Networks"

**Table S1**. Anchor atoms for reconstructing local reference frames of heavy atoms from 20 standard residue types

| Residue type | Atom type | Anchor atoms | | |
|---|---|---|---|---|
| | | A | B | C |
| ALA | CB | CA | N | CB |
| ARG | CB | CA | CB | CG |
| | CG | CB | CG | CD |
| | CD | CG | CD | NE |
| | NE | CD | NE | CZ |
| | CZ | NH1 | CZ | NH2 |
| | NH1 | NE | CZ | NH1 |
| | NH2 | NE | CZ | NH2 |
| ASN | CB | CA | CB | CG |
| | CG | OD1 | CG | ND2 |
| | ND2 | CG | CB | ND2 |
| | OD1 | CG | CB | OD1 |
| ASP | CB | CA | CB | CG |
| | CG | OD1 | CG | OD2 |
| | OD2 | CG | CB | OD2 |
| | OD1 | CG | CB | OD1 |
| CYS | CB | CA | CB | SG |
| | SG | CB | CA | SG |
| GLN | CB | CA | CB | CG |
| | CG | CB | CG | CD |
| | CD | OE1 | CD | NE2 |
| | OE1 | CD | CG | OE1 |
| | NE2 | CD | CG | NE2 |
| GLU | CB | CA | CB | CG |
| | CG | CB | CG | CD |
| | CD | OE1 | CD | OE2 |
| | OE1 | CD | CG | OE1 |
| | OE2 | CD | CG | OE2 |
| HIS | CB | CA | CB | CG |
| | CG | ND1 | CG | CD2 |
| | ND1 | CG | ND1 | CE1 |
| | CD2 | CG | CD2 | NE2 |
| | CE1 | ND1 | CE1 | NE2 |
| | NE2 | CD2 | NE2 | CE1 |

| Residue type | Atom type | Anchor atoms | | |
|---|---|---|---|---|
| | | A | B | C |
| ILE | CB | CA | CB | CG1 |
| | CG1 | CB | CG1 | CD1 |
| | CG2 | CB | CA | CG2 |
| | CD1 | CG1 | CB | CD1 |
| LEU | CB | CA | CB | CG |
| | CG | CD1 | CG | CD2 |
| | CD1 | CG | CB | CD1 |
| | CD2 | CG | CB | CD2 |
| LYS | CB | CA | CB | CG |
| | CG | CB | CG | CD |
| | CD | CG | CD | CE |
| | CE | CD | CE | NZ |
| | NZ | CE | CD | NZ |
| MET | CB | CA | CB | CG |
| | CG | CB | CG | SD |
| | SD | CG | SD | CE |
| | CE | SD | CG | CE |
| PHE | CB | CA | CB | CG |
| | CG | CD1 | CG | CD2 |
| | CD1 | CG | CD1 | CE1 |
| | CE1 | CD1 | CE1 | CZ |
| | CD2 | CG | CD2 | CE2 |
| | CE2 | CD1 | CE2 | CZ |
| | CZ | CE1 | CZ | CE2 |
| PRO | CB | CA | CB | CG |
| | CG | CB | CG | CD |
| | CD | CG | CD | N |
| SER | CB | CA | CB | OG |
| | OG | CB | CA | OG |
| THR | CB | CA | CB | OG1 |
| | OG1 | CB | CA | OG1 |
| | CG2 | CB | CA | CG2 |

| Residue type | Atom type | Anchor atoms | | |
|---|---|---|---|---|
| | | A | B | C |
| TRP | CB | CA | CB | CG |
| | CG | CD1 | CG | CD2 |
| | CD1 | CG | CD1 | NE1 |
| | CD2 | CE2 | CD2 | CE3 |
| | NE1 | CD1 | NE1 | CE2 |
| | CE2 | CD2 | CE2 | CZ2 |
| | CE3 | CD2 | CE3 | CZ3 |
| | CZ2 | CE2 | CZ2 | CH2 |
| | CZ3 | CE3 | CZ3 | CH2 |
| | CH2 | CZ2 | CH2 | CZ3 |
| TYR | CB | CA | CB | CG |
| | CG | CD1 | CG | CD2 |
| | CD1 | CG | CD1 | CE1 |
| | CE1 | CD1 | CE1 | CZ |
| | CD2 | CG | CD2 | CE2 |
| | CE2 | CD1 | CE2 | CZ |
| | CZ | CE1 | CZ | CE2 |
| | OH | CE2 | CZ | CE1 |
| VAL | CB | CA | CB | CG1 |
| | CG1 | CB | CA | CG1 |
| | CG2 | CB | CA | CG2 |
| all residues | N | Cprev[1] | N | CA |
| | CA | N | CA | C |
| | C | CA | C | Nnext[2] |
| | O | CA | O | Nnext |

[1] Backbone carbon of the preceding residue in the protein chain

[2] Backbone nitrogen of the next residue in the protein chain
